# Transcriptional regulation of the synthesis and secretion of farnesol in the fungus *Candida albicans:* Examination of the Homann transcription regulator knockout collection

**DOI:** 10.1101/2023.03.27.534401

**Authors:** Daniel J. Gutzmann, Jaxon J. Kramer, Brigid M. Toomey, Cory H. T. Boone, Audrey L. Atkin, Kenneth W. Nickerson

**Affiliations:** School of Biological Sciences University of Nebraska Lincoln, NE USA

**Keywords:** Candida albicans, Farnesol, Transcription factor mutant

## Abstract

*Candida albicans* is an efficient colonizer of human gastrointestinal tracts and skin and is an opportunistic pathogen. *C. albicans* exhibits morphological plasticity and the ability to switch between yeast and filamentous morphologies is associated with virulence. One regulator of this switch is the quorum sensing molecule farnesol which is produced by *C. albicans* throughout growth. However, the synthesis, secretion, regulation, and turnover of farnesol is not fully understood. To address this, we used our improved farnesol assay to screen a transcription regulator knockout library of 164 mutants for farnesol accumulation in whole cultures, pellets, and supernatants. All mutants produced farnesol and they averaged 9.2X more farnesol in the pellet than the supernatant. Nineteen mutants had significant differences with ten mutants producing more farnesol than their SN152^+^ parent while nine produced less. Seven mutants exhibited greater secretion of farnesol while two exhibited less. We examined the time course for farnesol accumulation in six mutants with the greatest accumulation differences and found that those differences persisted throughout growth and they were not time dependent. Significantly, two high accumulating mutants did not exhibit the decay in farnesol levels during stationary phase characteristic of wild type *C. albicans*, suggesting that a farnesol modification/degradation mechanism is absent in these mutants. Identifying these transcriptional regulators provides new insight into farnesol’s physiological functions regarding cell cycle progression, white-opaque switching, yeast-mycelial dimorphism, and response to cellular stress.

## Introduction

*Candida albicans* is an opportunistic pathogen that is present in most human gastrointestinal tracts and is an efficient colonizer of mucosal surfaces (Neville et al. 2015). A weakened immune system (HIV, chemotherapy, organ transplantation) or loss of competing flora (antibiotic treatment) allows for *C. albicans* to colonize and invade host tissues leading to candidiasis (Pfaller and Diekema 2007). Virulence of *C. albicans* is strongly linked to its ability to switch between yeast and filamentous morphologies (Romano 1966; Kadosh 2019) as mutants that have lost the ability to switch are avirulent (Lo et al. 1997; Saville et al. 2003). Dimorphism is also important for adaptation to different host environments (Noble et al. 2017; Alves et al. 2020) and evading host immune responses (Gow et al. 2012). One regulator of dimorphic switching is the quorum sensing molecule (QSM) farnesol (Hornby et al. 2001). Farnesol is produced by *C. albicans* cells continuously throughout growth (Boone et al. 2022), except in anaerobically grown cells (Dumitru et al. 2004) and opaque cells (Dumitru et al. 2007) where farnesol production is much lower. However, despite numerous publications on farnesol’s role as a QSM and virulence factor, the actual mechanisms regulating its synthesis and secretion remain unclear.

As an approach to identifying these genes and mechanisms, we tested the hypothesis that farnesol synthesis and secretion are regulated by the transcriptional networks involved in morphogenesis and virulence. The transcription regulator knockout library generated by Homann and colleagues has 165 different transcription regulators deleted, in most cases with two independently derived knockouts (Homann et al. 2009). Thus, we grew each of the library strains in liquid culture and then assayed them for total, extracellular, and intracellular farnesol by the improved methods of Boone et al. (2022). Key features of this assay include prevention of analyte loss by avoiding filtration and minimizing evaporation, while incorporating simultaneous cell lysis and analyte extraction by ethyl acetate. The assay enables comparison of whole culture values with the sum of their cell pellets and supernatants. Our results suggest farnesol accumulation is integrated with transcriptional regulators of cell cycle progression, white-opaque switching, yeast-mycelial dimorphism, and responses to cell stress. Our results provide new insights into the physiological role of farnesol and understanding of the regulatory mechanisms underlying its production.

## Materials and Methods

### Strains and media

The homozygous transcription regulator deletion mutants were obtained from the transcription regulator knockout library provided by Dr. Alexander Johnson’s lab (Homann et al. 2009) available through the Fungal Genomics Stock Center (Manhattan, Kansas, USA; McCluskey et al. 2010). The strains were maintained at 30°C on plates with YPD (1% yeast extract, 2% peptone, 2% glucose). The transcription regulator knockout mutants were constructed from *C. albicans* SN152, an auxotroph for arginine, histidine, and leucine. Each of the transcription regulator (TR) mutants is auxotrophic for arginine. A wild-type control strain was created by reintroduction of a single allele of HIS1 and LEU2 into the parent strain (Homann et al. 2009). Throughout this paper, we will refer to this control strain simply as SN152^+^. All assays with these mutants include the paired wild-type strain located in the H12 well of the X2 and Y2 independently derived deletion collections.

### Screening the transcription regulator deletion mutants for differences in accumulation farnesol secretion

The transcription regulator mutant collection was transferred by a 48-pin replicator to YPD plates and grown at 30°C for 48 hours. Mutants were then transferred to 3 mL YPD liquid medium and grown for 16 hours at 30°C with rotary agitation at 250 RPM. This culture was used as the inoculum (1:100) for 75 mL YPD in 250 mL flasks which were incubated 24 hours at 30°C, 250 RPM. Fifty mL of this culture was used for dry weight determination and 20 mL (2 X 10 mL) was used for determination of farnesol production as described by Boone et al. (2022). In this method, 10 mL was used to measure whole culture values while the other was centrifuged to obtain the pellet and supernatant values. Due to the large number of mutants in the collection, the mutants were assayed in batches of 10-20 mutants with the SN152^+^ control assayed in triplicate as part of each batch. Mutants with farnesol accumulation of greater than 1 log_2_ fold above or below the mean value for SN152^+^ were chosen for follow-up analysis. In follow-up analyses, E,E-farnesol accumulation was measured in at least three independent experiments using both the X and Y independently-derived TR deletion mutants (n=6).

### Temporal dynamics of farnesol accumulation

To examine the time course of farnesol production, we assayed the farnesol accumulation of the four highest and two lowest accumulating mutants at several cell densities throughout growth. Mutants were inoculated into to 6 mL YPD liquid medium and grown for 16 hours at 30°C with rotary agitation at 250 RPM. This culture was used as the inoculum (1:100) for replicate flasks for each time point with 75 mL YPD per mutant strain. Farnesol measurements were taken 12, 18, 24, 36, 48, and 80 hours post inoculation as describe by Boone et al. 2022. Two independent time courses were performed for the mutant strains and three independent time courses were performed for the SN152^+^ control. Farnesol accumulation values were normalized to the 50 mL dry weight at the indicated time point. To quantify differences in farnesol accumulation across time, area under the curve (AUC) analyses were performed in GraphPad Prism.

### Statistical analysis

Statistical analyses were performed using Microsoft Excel (Version 16.61, Microsoft Office, Las Vegas, NV, USA) and GraphPad Prism Software (Version 9.5.0, San Diego, CA, USA). All biological data are represented as mean ± SD of at least 3 biological replicates for both the X and Y independent mutants unless otherwise stated. Mutants with significant differences between the X and Y mutant were excluded and considered a false positive. Normality and homogeneity of variance were assessed by the D’ Agostino-Pearson omnibus (K2) test and Brown-Forsythe test, respectively. Differences between groups that were normally distributed and homoscedastic (equal variances) were assessed by One-way ANOVA with Dunnett’s multiple comparisons test. Differences between groups that were not normally distributed or heteroscedastic were assessed by the Kruskal-Wallis test with Dunn’s multiple comparisons tests. Both analyses were performed in GraphPad Prism. Differences were considered significant at p<0.05 (*p<0.05, **p<0.01, ***p<0.001).

## Results

### General design and reproducibility of the screen for farnesol regulators

Homann et al. (2009) screened their collection of 165 independent transcription regulator mutants for possible differences in 55 growth conditions. It was a monumental piece of work. However, for practical reasons they limited themselves to phenotypes discernible via colonies on agar plates. We have now extended the list of phenotypes to include accumulation of both intracellular and extracellular farnesol as determined by our improved GC assay (Boone et al, 2022). For the initial screen, 164 mutants were screened in 12 batches. One mutant *ΔΔhfl1* [orf19.3063] was not screened as it exhibited a severe growth defect on YPD. Each batch included duplicate or triplicate SN152^+^ controls, for a total of 198 cultures and 594 GC-FID runs. Farnesol accumulation values were normalized on a per cell basis using the 50 mL dry weight value of the culture assayed. In this initial screen, the paired WT controls were run a total of 34 times and their whole culture mean E,E-farnesol accumulation value was 5.29 ± 1.12 ng/µL/ 50 mL dry weight (8.27 ± 2.8 μM).

For the rescreen, 27 strains were selected because their farnesol values differed from the average values for SN152^+^ by a log_2_ fold change of > 1.0. To these we added 3 strains (*ΔΔupc2, ΔΔtac1, and ΔΔcsr1*) because prior reports from the literature indicated they were likely to have altered farnesol accumulations. These 30 strains were then rescreened five more times, two from the X plate and three from the Y plate (total n =6) in 10 batches including the SN152^+^ parent in biological duplicate or triplicate, requiring 180 flasks and 540 GC measurements. Thus the rescreen data set examines 30 transcriptional regulator mutants (n=6), reporting the means ± SD for normalized farnesol accumulation as detected in whole cultures, pellets, and supernatants as well as the supernatant/pellet ratio. Composite data sets for the initial screen (File S1), rescreen (File S2) and farnesol growth curves (File S3) are available at Figshare. We now describe several highlights from these data sets.

### Mutants altered in farnesol accumulation

Our initial screen of 164 strains, all from the X plate, was done with an n = 1 comparing whole culture values versus the sum of the cell pellet and supernatant values. The 164 TR mutants were assayed in 12 batches which always included a biological duplicate or triplicate of the SN152^+^ parent. The whole culture accumulation and supernatant/pellet ratios are shown in Fig. 1. The distribution of the 34 values for SN152^+^ (red) is shown in Fig. 1 where they can be seen in comparison with the distribution of the 164 TR mutants (black). As expected (Boone et al. 2022), the whole culture (W) and pellet (P) + supernatant (S) values were always very close to one another, with the average WPS relative error values (W-(P+S)/W) being 0.07 ± 0.06 for E,E-farnesol.

**Fig. 1.**
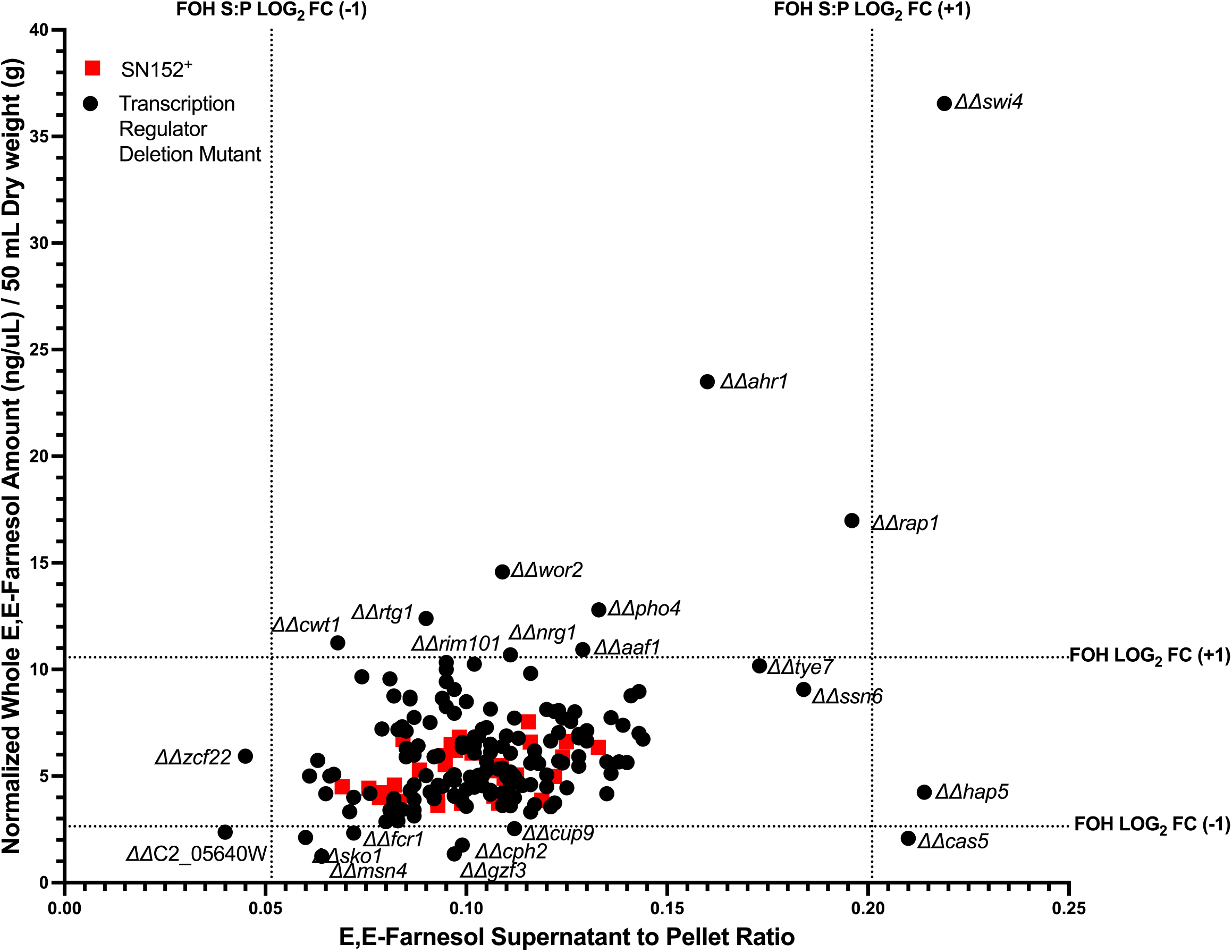
Initial screen E,E-farnesol supernatant to pellet ratio vs E,E-farnesol whole culture accumulation at 24 hours post inoculation. Data represents the accumulation values for each of 164 transcription regulator mutants (black) (n=1) and each SN152^+^ replicate (red) (n=34). The red X indicates the mean value for the SN152^+^ replicates (n=34). Dotted lines represent a log2 fold change increase or decrease of 1 for farnesol from the mean for SN152^+^.

The vast majority of the 164 TR mutants were unchanged from the SN152^+^ replicates (Fig. 1). We then identified all mutants which were > 1 log_2_ above or below the mean values, as indicated by the 4 dotted lines in Fig. 1, and designated them as mutants of interest to be rescreened with greater accuracy. There were 27 mutants with higher or lower farnesol accumulation. Six mutants (*ΔΔcwt1, ΔΔrim101, ΔΔcrz2, ΔΔzcf1,ΔΔtea1, ΔΔrme1*) were designated as false positives because their rescreened values from the X and Y plates differed significantly or their rescreen values differed significantly from the initial screen. Thus, we identified ten mutants with elevated farnesol (Fig. 2A), nine with reduced farnesol (Fig. 2B), seven with elevated supernatant/pellet ratios (Fig. 2C), and two with lowered farnesol supernatant/pellet ratios (Fig. 2C). Each of these mutants is described in greater detail in Table 1.

**Fig. 2.**
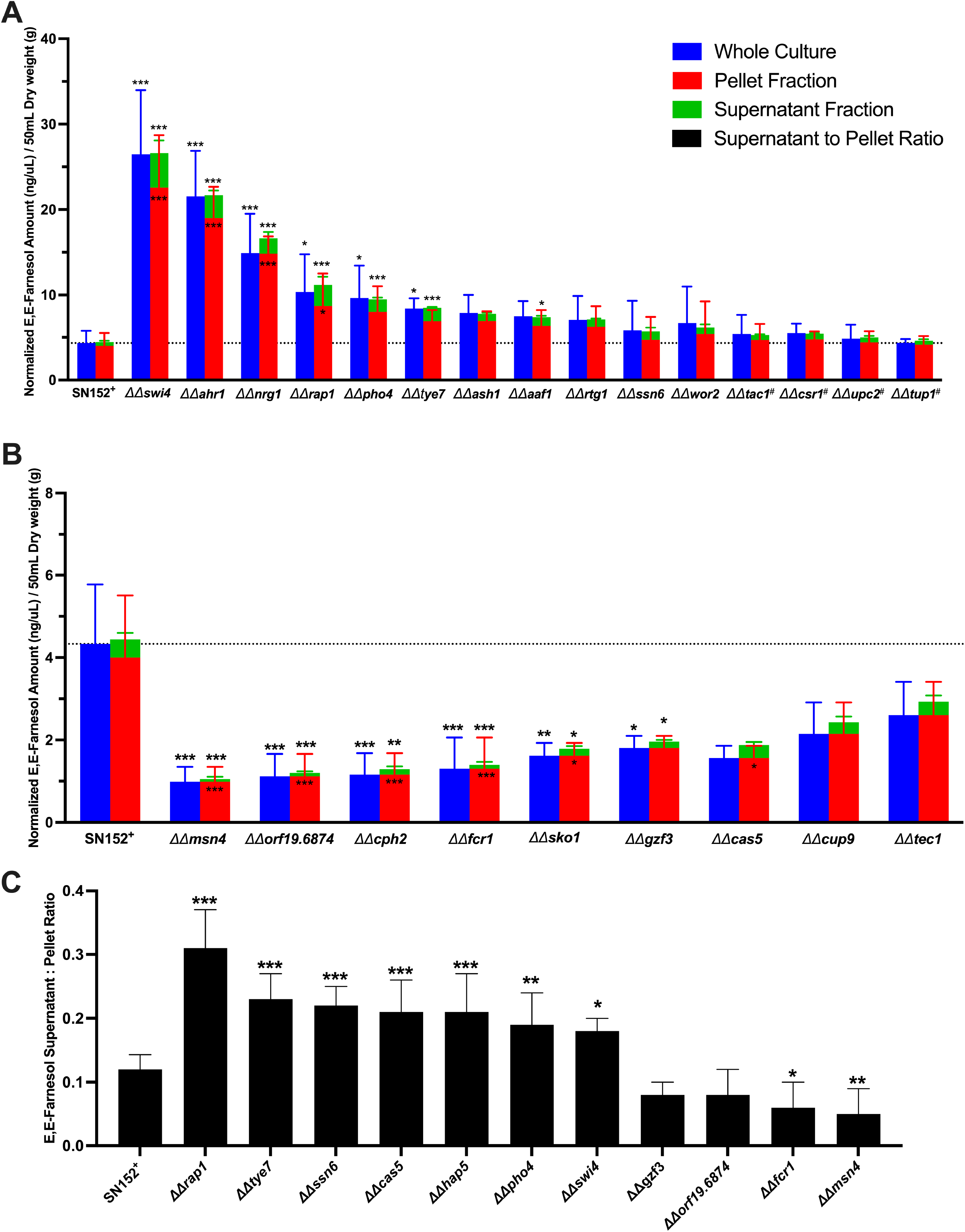
Transcription regulator mutants differing in farnesol accumulation and localization. Accumulation values were measured 24 hours post inoculation at 30°C in YPD for each transcription regulator mutant in both the X (3) and Y (3) independent mutants (total n=6) and the SN152^+^ parent (n=22 batches). Data represents the mean accumulation value ± SD (A,B) or mean farnesol supernatant to pellet ratio ± SD The mutants added because of literature precedent are indicated by #. Differences between groups that were normally distributed and that had equal variances (C) were accessed by One-way ANOVA with Dunnett’s multiple comparisons test. Differences between groups that were not normally distributed or heteroscedastic (A,B) were accessed by Kruskal-Wallis test with Dunn’s multiple comparisons tests. Differences were considered significant at p<0.05 (*p<0.05, **p<0.01, ***p<0.001).

**Table 1.** Summary of transcription regulator mutants with statistically significant differences in farnesol accumulation in YPD media at 30°*C.*^a/^

| Mutant | *S. cerevisiae* ortholog or best hit from CGD | Mean whole culture E,E-farnesol log<sub>2</sub> fold change vs WT control ± SD (n=6) | Mean supernatant fraction E,E-farnesol log<sub>2</sub> fold change vs WT control ± SD (n=6) | Predicted or known biological process(es) in *Candida albicans* adapted from CGD (Skrzpek et al. 2017) |
| --- | --- | --- | --- | --- |
| E,E-farnesol over accumulating mutants |  |  |  |  |
| *ΔΔswi4* | *SWI4* | 2.57*** ± 0.39 | 3.11*** ± 0.50 | Component of SBF complex, G1/S progression |
| *ΔΔahr1* | *LYS14* | 2.27*** ± 0.42 | 2.56 *** ± 0.32 | Regulation of adhesion genes, white/opaque switch, repressor of START |
| *ΔΔnrg1* | *NRG2* | 1.73*** ± 0.45 | 1.93*** ± 0.49 | Regulator of hyphal gene induction, virulence, chlamydospore formation, stress response |
| *ΔΔrap1* | *RAP1* | 1.15*** ± 0.62 | 2.42*** ± 0.48 | Telomere maintenance |
| *ΔΔpho4* | *PHO4* | 1.07*** ± 0.53 | 1.68*** ± 0.23 | Phosphate acquisition |
| *ΔΔtye7* | *TYE7* | 0.95** ± 0.21 | 1.83*** ± 0.07 | Regulator of glycolysis and carbohydrate metabolism |
| *ΔΔash1* | *ASH1* | 0.82* ± 0.43 | 0.92** ± 0.28 | GATA-like transcription factor required for WT virulence and filamentous growth |
| *ΔΔaaf1* | *none* | 0.76* ± 0.31 | 1.21*** ± 0.19 | Expression in *S. cerevisiae* increases adherence |
| *ΔΔrtg1* | *RTG1* | 0.63 ± 0.49 | 0.96** ± 0.20 | Regulation of sphingolipid homeostasis, galactose catabolism |
| $\Delta\Delta ssn6$ | *CYC8* | $0.20 \pm 0.90$ | $0.99^{***} \pm 0.73$ | Forms a corepressor complex with Tup1, regulates a distinct set of genes different than Tup1 in *C. albicans* |
| $\Delta\Delta wor2$ | *UME6* | $0.43 \pm 0.82$<br>(p=0.999) | $0.59 \pm 0.83$<br>(p=0.217) | Regulator of white/opaque switching, maintenance of opaque state |
| $\Delta\Delta tac1^{\#}$ | *HAL9* | $0.23 \pm 0.56$<br>(p=0.999) | $0.36 \pm 0.26$<br>(p=0.919) | Transcriptional activator of drug-responsive genes *CDR1* and *CDR2* |
| $\Delta\Delta csr1^{\#}$ | *ZAP1* | $0.32 \pm 0.31$<br>(p=0.984) | $0.60 \pm 0.13$<br>(p=0.195) | Transcription factor, role in zinc homeostasis |
| $\Delta\Delta upc2^{\#}$ | *UPC2* | $0.11 \pm 0.43$<br>(p=0.999) | $0.31 \pm 0.46$<br>(p=0.980) | Regulator of ergosterol biosynthetic genes and sterol uptake |
| $\Delta\Delta tup1^{\#}$ | *TUP1* | $0.0 \pm 0.15$<br>(p=0.999) | $-0.02 \pm 0.47$<br>(p=0.999) | Transcriptional corepressor with *Ssn6p*, regulates morphological switching, farnesol response, germ tube formation |
| E,E-farnesol under accumulating mutants |  |  |  |  |
| $\Delta\Delta$<br>*orf19.6874*<br>(*C2_05640*<br>*W_A*) | *BAS1* | $-2.10^{***} \pm 0.73$ | $-2.65^{***} \pm 1.03$ | Transcription factor, null mutant exhibits increase in invasive growth |
| $\Delta\Delta msn4$ | *MSN2* | $-1.89^{***} \pm 0.37$ | $-2.40^{***} \pm 0.65$ | Zinc finger transcription factor, not a significant stress response regulator in *C. albicans* |
| $\Delta\Delta cph2$ | *HMS1* | $-1.93^{***} \pm 0.63$ | $-2.04^{***} \pm 0.91$ | Transcriptional activator of hyphal growth |
| $\Delta\Delta fcr1$ | *CAT8* | $-1.97^{***} \pm 0.88$ | $-3.68^{***} \pm 3.83$ | Negative regulator of fluconazole/ketoconazole/brefeldin A resistance |
| $\Delta\Delta sko1$ | *SKO1* | $-1.93^{***} \pm 0.39$ | $-1.41^{***} \pm 0.46$ | Cell wall damage response, morphological switching, oxidative stress response |
| $\Delta\Delta gzf3$ | *GZF3* | $-1.32^{***} \pm 0.32$ | $-1.63^{***} \pm 0.54$ | GATA type transcription factor, ortholog regulates nitrogen catabolic gene expression |
| $\Delta\Delta cas5$ | *MOT3* | $-1.16^{**} \pm 0.13$ | $-0.51 \pm 0.32$ | Cell wall homeostasis, adherence, stress response, cell cycle |
| $\Delta\Delta cup9$ | *TOS8* | $-1.08^{**} \pm 0.63$ | $-0.85 \pm 0.86$ | Represses SOK1 expression in response to farnesol inhibition |
| $\Delta\Delta tec1$ | *TEC1* | $-0.75^{*} \pm 0.66$ | $-0.58 \pm 0.76$ | Hyphal gene regulator, white cell pheromone response |

a/ Log<sub>2</sub> fold change production values ± SD represent the mean farnesol accumulation value of 3 X and 3 Y (n=6) independently derived mutant compared to the mean farnesol accumulation value of the paired SN152 parent (n=22). Both over and under accumulating mutants are ordered by how much they exceed the SN152 parent. The four mutants added because of literature precedent are indicated by #. Differences between mutants (n=6) and the SN152<sup>+</sup> parent (n=22 batches) were accessed by One-way ANOVA with Dunnett's multiple comparisons test using GraphPad Prism. Differences were considered significant at p<0.05 (\*p<0.05, \*\*p<0.01, \*\*\*p<0.001).

Of the 19 mutants with statistically significant differences in farnesol accumulation, only *ΔΔnrg1* had been identified in prior literature (Kebaara et al. 2008). Ten mutants (*ΔΔswi4, ΔΔahr1, ΔΔnrg1, ΔΔrap1, ΔΔpho4, ΔΔtye7, ΔΔaaf1, ΔΔash1, ΔΔrtg1, ΔΔssn6*) had significantly increased farnesol accumulation either in the whole culture or supernatant fraction compared to the SN152^+^ parent strain (Table 1). The two highest accumulators (*ΔΔswi4* and *ΔΔahr1)* have known roles in cell cycle progression (Hussein et al. 2011; Sellam et al. 2019). Additionally, two farnesol over accumulating mutants (*ΔΔtye7* and *ΔΔrtg1*) have roles in carbohydrate metabolism (Askew et al. 2011; Moreno-Velásquez et al. 2020), four (*ΔΔahr1, ΔΔnrg1, ΔΔash1, ΔΔssn6*) have a role in dimorphism (Fu et al. 1998; Murad et al. 2001; Inglis and Johnson 2002; García-Sánchez et al. 2005), and two (*ΔΔahr1, ΔΔssn6)* have roles in white/opaque switching (Zordan et al. 2007; Wang et al. 2011; Hernday et al. 2016). *ΔΔwor2* appeared to have a modest increase in accumulation but this did not reach the significance threshold (Table 1, Fig. 2A). We note that another regulator of white/opaque switch, Czf1, has role in coordinating switching with the response to farnesol (Langford et al. 2013).

Nine mutants (*ΔΔorf19.6874, ΔΔmsn4, ΔΔcph2, ΔΔfcr1, ΔΔsko1, ΔΔgzf3, ΔΔcas5, ΔΔcup9, ΔΔtec1*) had significantly decreased farnesol accumulation either in the whole culture or supernatant fraction compared to the SN152^+^ parent strain (Table 1). Three of these low farnesol accumulating mutants (*ΔΔmsn4, ΔΔsko1, ΔΔcas5*) have known roles in stress response to cell wall damage and oxidative stress(Nicholls et al. 2004; Bruno et al. 2006; Rauceo et al. 2008; Xie et al. 2017) and one mutant (*ΔΔcup9*) has a known role in farnesol response by mediating Nrg1 degradation in the absence of farnesol (Lu et al. 2014).

The three mutants chosen because of literature precedent had elevated farnesol accumulation but did not reach statistical significance (*ΔΔcsr1, ΔΔupc2, ΔΔtac1*) (Table 1). Ganguly et al. (2011) reported decreased accumulation of farnesol by the *ΔΔcsr1* mutant. Notably, the assay conditions used in our screen (YPD, 30°C) differ significantly from those used in prior studies of *ΔΔcsr1* which employed biofilm forming conditions at 37°C in SPIDER medium (Ganguly et al. 2011). Thus, the dynamics of farnesol accumulation are likely influenced not only by temperature, but also by growth medium (Boone et al. 2022).

### Mutants altered in the farnesol supernatant/pellet ratio

The supernatant/pellet (S/P) ratios for farnesol were determined for each mutant (n=1) in comparison to those for their SN152^+^ control (Fig. 1). The value for SN152^+^ (n= 22 batches) was 0.12 ± 0.02 for farnesol. As we had observed previously (Boone et al. 2022), farnesol was predominantly retained in the cell pellet. The S/P ratios were all tightly clustered from 0.07-0.14 except for four mutants (*ΔΔmsn4,ΔΔfcr1, ΔΔgzf3, and ΔΔorf19.6874*) which exhibited less secretion (0.05-0.08) and seven (*ΔΔswi4, ΔΔpho4, ΔΔcas5, ΔΔhap5, ΔΔssn6, ΔΔtye7, and ΔΔrap1*) which exhibited greater secretion (0.19-0.30) (Fig. 2C). Inexplicably, *ΔΔzcf22* was not among the rescreened mutants; it should likely be included among the mutants which exhibited less secretion.

Two mutants, *ΔΔhap5* and *ΔΔcas5* are of special interest in that they exhibit greater secretion without being accompanied by greater total farnesol production (Fig. 1). Thus, they target secretion specifically. Mutants such as *ΔΔswi4, ΔΔahr1, and ΔΔrap1* with much greater farnesol production (Fig. 2A, Table 1) could exhibit a higher supernatant to pellet ratio because internal and external farnesol are in equilibrium or there is a need to keep internal farnesol levels below a critical threshold value. In both cases, the greater secretion would be an indirect consequence of greater production.

### Temporal dynamics of farnesol accumulation

The initial screen and rescreen for farnesol accumulation were done at a single time point 24 hours post inoculation. To verify that farnesol differences we observed were not time dependent, we extended the assay to 12, 18, 24, 36, 48, and 80 hours post inoculation for four over accumulating (*ΔΔswi4, ΔΔahr1, ΔΔrap1, ΔΔnrg1*) and two under accumulating (*ΔΔmsn4, ΔΔ19.6874*) mutants (Figs. 3 and 4, respectively), with n=2 for each mutant and n=3 for the SN152+ control. Growth of the six mutant strains paralleled the growth of SN152^+^ (Figs. 3A, 4A). As expected, the four over accumulating mutants had elevated farnesol at all time points in both the whole culture (Fig. 3B) and supernatant fractions (Fig. 3D) while the two under accumulating mutants had decreased farnesol in both the whole culture (Fig. 4B) and supernatant fractions (Fig. 4D). To compare total farnesol accumulations, area under the curve analyses (AUC) were performed. The four over accumulating mutants *ΔΔswi4, ΔΔrap1, ΔΔahr1,* and *ΔΔnrg1* had significantly higher AUC in the whole culture (Fig. 3C) and supernatant (Fig. 3E) than the SN152^+^ parent while *ΔΔorf19.6874* and *ΔΔmsn4* had significantly lower supernatant AUC (Fig. 4E). Of great interest, two of the over accumulating mutants (*ΔΔswi4, ΔΔrap1)* still had significant farnesol remaining at 80 hours (Figs. 3B, 3D). They did not exhibit the decay in farnesol levels during stationary phase characteristic of wild type *C. albicans* (Boone et al. 2022) as well as SN152^+^ and the other two over accumulating mutants tested (Fig. 3B), suggesting that a farnesol modification or degradation mechanism is absent from these two mutants.

**Fig. 3.**
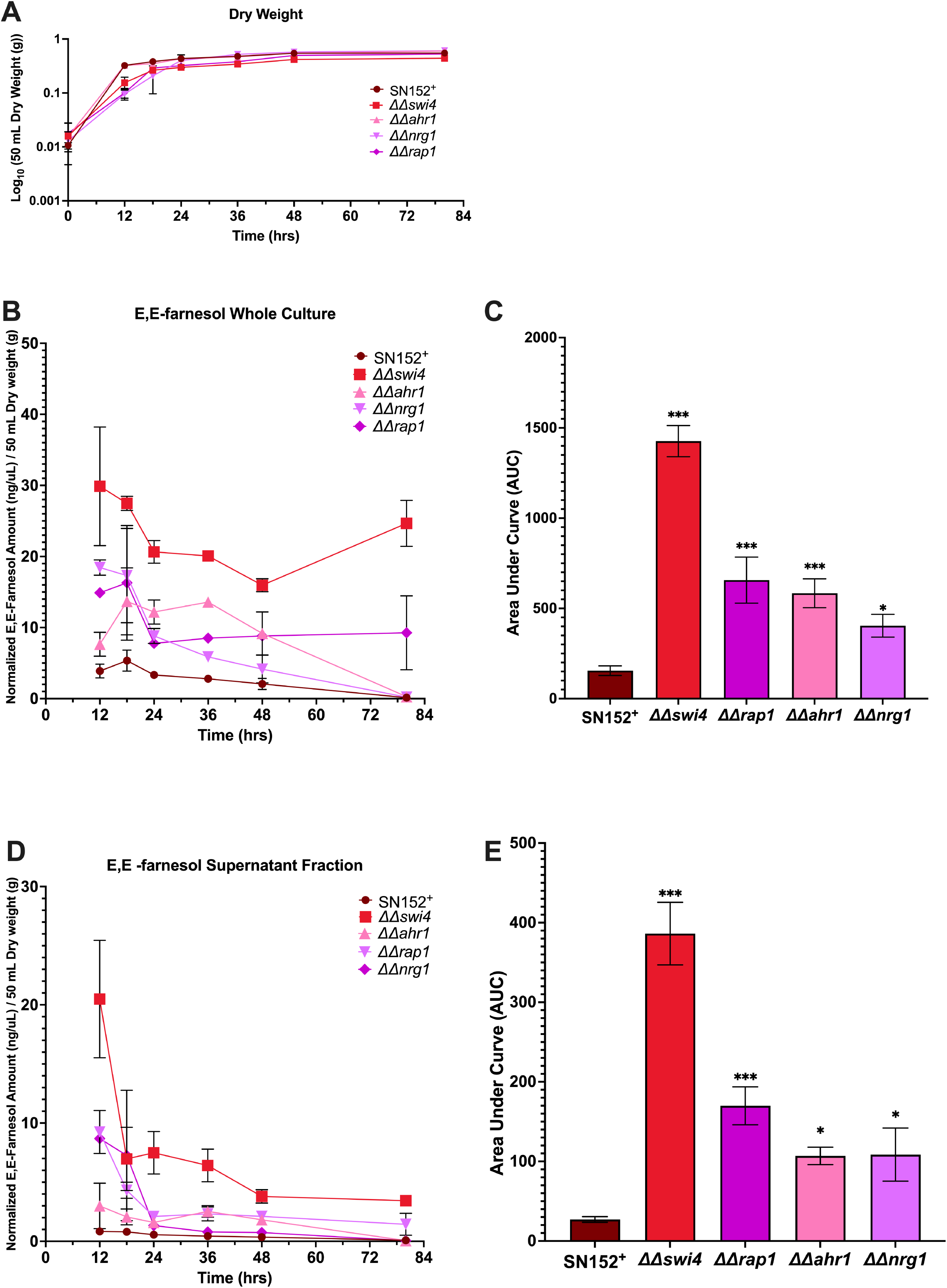
Temporal dynamics of farnesol accumulation in *ΔΔswi4, ΔΔahr1, ΔΔrap1, and ΔΔnrg1.* Farnesol accumulation was accessed at 12, 18, 24, 36, 48, 80 hours post inoculation at 30°C in YPD for each transcription regulator mutant in two independent growth curves (n=2) and 3 independent growth curves for SN152^+^ (n=3). Dry weights were accessed as each time point and are presented as the mean dry weight ± SEM (A). Farnesol accumulation data represents the mean dry weight normalized farnesol accumulation value ± SEM in either the whole culture (B) or the supernatant fraction (D). To access overall farnesol production, area under the curve analyses (AUC) were performed and data are the mean AUC ± SE for the whole culture (C) or supernatant fraction (E). Differences in the AUC between groups were accessed by One-way ANOVA with Dunnett’s multiple comparisons test. Differences were considered significant at p<0.05 (*p<0.05, **p<0.01, ***p<0.001).

**Fig. 4.**
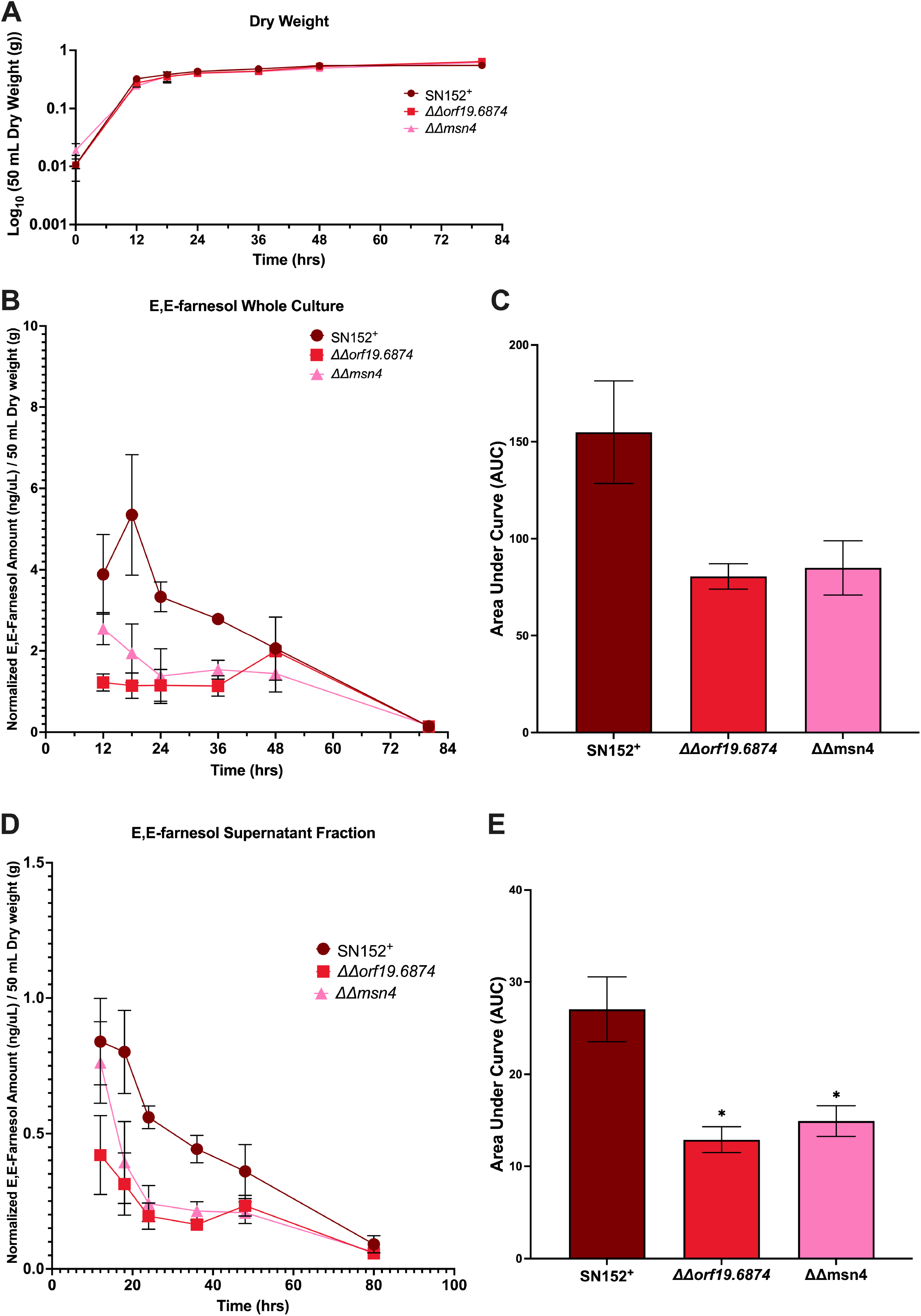
Temporal dynamics of farnesol accumulation in *ΔΔorf19.6874, and ΔΔmsn4.* Farnesol accumulation was accessed at 12, 18, 24, 36, 48, 80 hours post inoculation at 30°C in YPD for each transcription regulator mutant in two independent growth curves (n=2) and 3 independent growth curves for SN152^+^ (n=3). Dry weights were accessed as each time point and are presented as the mean dry weight ± SEM (A). Farnesol accumulation data represents the mean dry weight normalized farnesol accumulation value ± SEM in either the whole culture (B) or the supernatant fraction (D). To access overall farnesol production, area under the curve analyses (AUC) were performed and data are the mean AUC ± SE for the whole culture (C) or supernatant fraction (E). Differences in the AUC between groups were accessed by One-way ANOVA with Dunnett’s multiple comparisons test. Differences were considered significant at p<0.05 (*p<0.05, **p<0.01, ***p<0.001).

### Opaque phenotype of *ΔΔtup1*

One surprising result of the screen (Fig. 1) was that *ΔΔtup1* did not appear to over accumulate farnesol in these assay conditions (Table 1). This result was surprising because we had previously reported that the *ΔΔtup1* mutant produced 17 times more farnesol than its parent (Kebaara et al. 2008). A possible explanation for this discrepancy is that *C. albicans* can undergo phenotypic switching between two heritable states: white and opaque, each of which is normally stable for thousands of cell divisions. Switching between the two cell types is reversible and occurs without any chromosomal rearrangements or sequence changes. Critically, Tup1 is a key repressor of the opaque state and Alkafeef et al. (2018) have shown that loss of *TUP1* is sufficient to induce the opaque phase, even in a MTL a/α background (Alkafeef et al. 2018). The pressure of *ΔΔtup1* to spontaneously convert to the opaque phase is strong (Alkafeef et al. 2018) and we already know that wild type opaque cells usually produce far less farnesol than do white cells (Dumitru et al. 2007). Given this, we investigated whether the *ΔΔtup1* mutant in the Homann collection is in the opaque phase under the assay conditions performed in the screen (YPD, 30°*C*). It is important to note that *ΔΔtup1* cells are locked in the filamentous morphology (Kebaara et al. 2008), making visual determination between white and opaque cells more difficult. When plated on YPD medium supplemented with phloxine B and incubated at 30°C, the *ΔΔtup1* colonies were pink while the SN152^+^ colonies plated under the same conditions were white (Fig. 5A). The pink colonies for *ΔΔtup1* are consistant with the cells being in the opaque phase. This view was supported by microscopic examination of the *ΔΔtup1* cells which showed numerous short but elongated cells consistent with the opaque state (Fig. 5B). The typical budding yeast cells of SN152^+^ are included for comparison (Fig. 5B). We are currently investigating the conditions that influence the farnesol production by white and opaque *ΔΔtup1* cells.

**Fig. 5.**
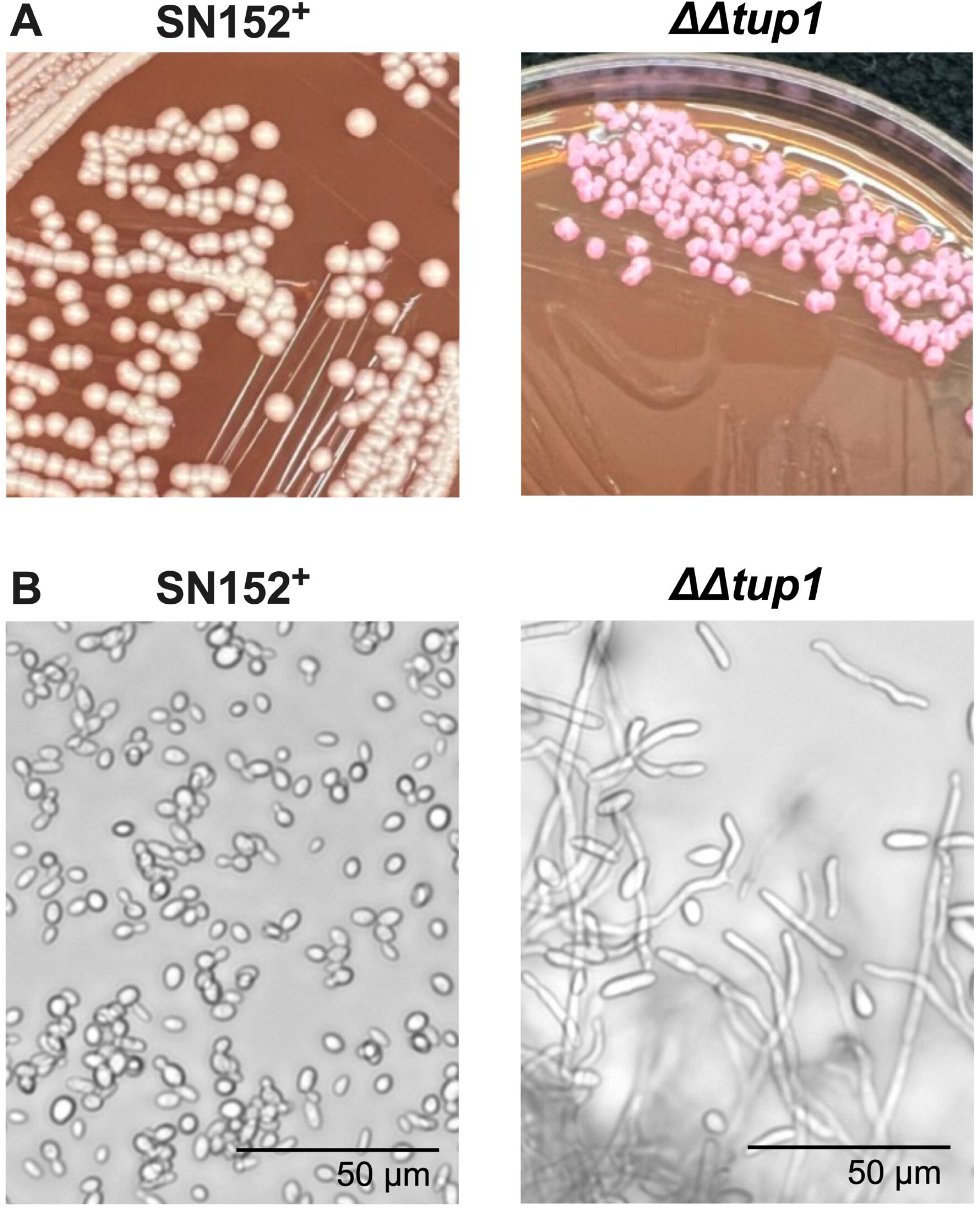
Characterization of *ΔΔtup1* morphology in conditions used for the screen. (A) Representative micrographs of the colony morphology of SN152^+^ (white) and *ΔΔtup1* (pink) on YPD agar supplemented with phloxine B grown at 30°C. (B) Representative micrographs of cellular morphology of SN152^+^ (yeast) and *ΔΔtup1* (hyphae and elongated cells).

## Discussion

We examined 164 independent transcriptional regulator knockout mutants for their accumulation and secretion of farnesol. This project was undertaken to define the genes needed for farnesol’s synthesis, secretion, and regulation. For logistical reasons, the initial screen and rescreen cultures were grown at 30°C in liquid YPD medium and sampled after 24 hours when the cells were in early stationary phase. The 24 hour time point was chosen based on our prior measurements with wild type *C. albicans* SC5314 over 72 hours of culture which showed that farnesol accumulation generally followed increased cell mass, peaked in early stationary phase, and then declined sharply after ∼ 30 hours (Boone et al. 2022). Thus, the 24 hour time point was intended to cover the peak of farnesol accumulation. Similarly, our choice of YPD as a growth medium was to insure that all of the TR mutants grew well by 24hrs and indeed 164 out of 165 had done so.

With the advantage of now having reliable measures for farnesol presence in whole cultures, cell pellets, and supernatants, we can approach the genetic basis for their synthesis, secretion, regulation, and turnover. As expected, most of the TR mutants were unchanged for these parameters (Fig. 1) while a few potentially useful mutants gave significantly higher and lower levels of farnesol. Significantly, no mutants were entirely devoid of farnesol; its levels often varied but they never dropped to zero. A tentative conclusion from this observation is that farnesol has an essential physiological function over and above acting as a quorum sensing molecule and virulence factor or that it is the product of a redundant biosynthetic pathway. In examing mutants with significant differences in farnesol production, several patterns in the transcriptional networks emerge that may highlight farnesol’s physiological functions. These networks include: 1) white-opaque switching, 2) yeast-mycelia dimorphism, 3) response to cell stress, and 4) cell cycle progression. These networks integrate environmental signals to influence the morphology of C*. albicans*. Firstly, three of the transcription regulators studied, Ahr1, Ssn6, and Tup1, have crucial roles in white-opaque switching (Hernday et al. 2013; Hernday et al. 2016; Alkafeef et al. 2018). Given that farnesol is toxic to opaque cells (Dumitru et al. 2007), it is intriguing that the *AHR1* deletion mutant over accumulates farnesol while also promoting a higher frequency of white-opaque switching (Hernday et al. 2013). Additionally, it is unclear whether Ssn6 and Ahr1 participate in the response to farnesol, but there is ample precedent given that another transcriptional regulator of the white-opaque switch, Czf1, is involved in farnesol response. Secondly, in addition to Nrg1 and Tup1 (Kebaara et al. 2008), four more transcriptional regulators of yeast-mycelia dimorphism have been demonstrated in this study to influence farnesol accumulation including Cph2, Ash1, Ssn6, and Tec1. Thirdly, *C. albicans* is especially resistant to farnesol compared to other fungi (Semighini et al. 2006; Brasch et al. 2013; Wang et al. 2014), but the mechanism of this tolerance is elusive. One possibility concerns the protective effect of using longer chain ubiquinones (UQ). *C. albicans* and *Candida dubliniensis* use UQ9 rather than UQ7 as in other *Candida* species or UQ6 as in *S. cerevisiae* (Pathirana et al. 2020). Another possibility could be due to differences in their response to cellular stress. Given that Msn4 and Cas5 have roles in stress response (Nicholls et al. 2004; Xie et al. 2017) and influence farnesol accumulation and secretion (Fig. 2), it is likely they may also contribute to farnesol tolerance. Fourthly, cell cycle arrest can also induce filamentous growth of *C. albicans,* either through depletion of cyclins such as Cln3, Clb2, or Clb3 (Bachewich et al. 2005; Bensen et al. 2005), depletion of polo-like kinase Cdc35 (Bachewich et al. 2003), or through treatment with cell cycle inhibitors like hydroxyurea (Bachewich et al. 2005; Chen et al. 2018). Interestingly, expression of the G1/S cyclins *PCL2*, *CLN3*, and *HGC1* are farnesol regulated (Enjalbert and Whiteway 2005). Given that *PLC2* and *CLN3* are Swi4 regulated (Hussein et al. 2011), how these cyclins influence farnesol accumulation is an important question for subsequent studies.

There are many questions which can now be approached via the genetic information gleaned from this screen. Among them are: 1/ How is carbon flow through the farnesyl pyrophosphate (FPP) branch point regulated? 2/ Is farnesol made directly from FPP by an appropriate pyrophosphatase (e.g. Dpp1p, Dpp2p, or Dpp3p) or are there other more easily regulated and/or safer biosynthetic pathways? A pathway suggested from *Arabidopsis* invokes the necessary turnover of farnesylated proteins, where proteolytic degradation of those proteins releases farnesylated cysteine which can be cleaved by a ligase to release farnesaldehyde which is then converted to farnesol by a farnesol dehydrogenase (Bhandari et al. 2010). 3/ Is the disappearance of farnesol later in stationary phase due strictly to evaporation of a volatile molecule or to enzymatic conversion of farnesol to, for instance, 2,3-dihydrofarnesol (Brasch et al. 2013; Costa et al. 2020). Significantly, regarding its ability to block germ tube formation and hyphal development, 2,3-dihydrofarnesol is inactive; with only 0.34% of the activity of farnesol (Shchepin et al. 2003). This question has now been partially resolved by extending the *ΔΔswi4* and *ΔΔrap1* mutant analyses to 80 hours post inoculation (Fig. 3B). While wild type SC5314 (Boone et al. 2022), SN152^+^, and most of the mutants examined had little farnesol remaining by 80 hours, *ΔΔswi4* and *ΔΔrap1* differed in that they maintained high levels of farnesol. Clearly, something other than evaporation is occurring. There is ample precedent for farnesol modifying enzymes in fungi. For instance, trans-trans farnesol can be converted to a cis-trans farnesol by *Helminthosporium sativum* (Imai and Marumo 1974). Presumably, the corresponding enzymes for *C. albicans* are not made by *Δswi4* and *ΔΔrap1,* and we are currently using these genetic clues to follow the metabolic disappearance of farnesol during stationary phase.

## Author contributions

DJG: conceptualization, methodology, validation, investigation, writing — original draft, writing — review and editing, visualization, formal analysis, data curation, project administration; JJK: validation, investigation, writing — review and editing; BMT: validation, investigation, writing — review and editing; CHTB: methodology, validation, writing – review and editing; ALA: conceptualization, methodology, resources, supervision, writing — review and editing, project administration; KWN: conceptualization, methodology, resources, supervision, writing — original draft, writing — review and editing, project administration, funding acquisition.

## Data availability

The datasets generated during and/ or analyzed during the current study are available at FigShare. This includes the initial screen (File S1), rescreen dataset (File S2), and farnesol growth curves (File S3).

## Acknowledgments

We are grateful to the Fungal Genetics Stock Center (FGSC) for providing strains, to Prof. Rebecca Roston and Zach Shomo for use and assistance with the gas chromatography system, and to Prof. Haoping Liu for pointing out the possible role of opaque *ΔΔtup1*.

## Compliance with ethical standards

### Funding

This work was supported by Ann L. Kelsall and the Farnesol and *Candida albicans* Research Fund, University of Nebraska Foundation. This research did not receive any specific grant from funding agencies in the public, commercial, or not-for-profit sectors.

### Ethical approval

This article does not contain any studies with human participants or animals performed by any of the authors.

